# E-Cannula reveals anatomical diversity in sharp-wave ripples as a driver for the recruitment of distinct hippocampal assemblies

**DOI:** 10.1101/2021.10.22.465514

**Authors:** Xin Liu, Satoshi Terada, Jeong-Hoon Kim, Yichen Lu, Mehrdad Ramezani, Andres Grosmark, Attila Losonczy, Duygu Kuzum

## Abstract

The hippocampus plays a critical role in spatial navigation and episodic memory. However, research on *in vivo* hippocampal activity dynamics has mostly relied on single modalities such as electrical recordings or optical imaging, with respectively limited spatial and temporal resolution. This technical difficulty greatly impedes multi-level investigations into network state-related changes in cellular activity. To overcome these limitations, we developed the E-Cannula integrating fully transparent graphene microelectrodes with imaging-cannula. The E-Cannula enables the simultaneous electrical recording and two-photon calcium imaging from the exact same population of neurons across an anatomically extended region of the mouse hippocampal CA1 stably across several days. These large-scale simultaneous optical and electrical recordings showed that local hippocampal sharp wave ripples (SWRs) are associated with synchronous calcium events involving large neural populations in CA1. We show that SWRs exhibit spatiotemporal wave patterns along multiple axes in 2D space with different spatial extents (local or global) and temporal propagation modes (stationary or travelling). Notably, distinct SWR wave patterns were associated with, and decoded from, the selective recruitment of orthogonal CA1 cell assemblies. These results suggest that the diversity in the anatomical progression of SWRs may serve as a mechanism for the selective activation of the unique hippocampal cell assemblies extensively implicated in the encoding of distinct memories. Through these results we demonstrate the utility of the E-Cannula as a versatile neurotechnology with the potential for future integration with other optical components such as green lenses, fibers or prisms enabling the multi-modal investigation of cross-time scale population-level neural dynamics across brain regions.

## Introduction

Large-scale electrophysiological and optical recordings have significantly advanced our understanding of neural circuit dynamics in the mammalian neocortex. However, similar experiments and analyses remain challenging in circuits that are located deep in the brain, such as the hippocampus, a region critical for spatial navigation and episodic memory^1,2^. To date, most of the knowledge about hippocampal neural dynamics comes from experimental studies tracking responses from a small number of neurons monitored by conventional microelectrodes. However, understanding hippocampal functions requires monitoring the activity of large populations of hippocampal cells while recording the network level oscillations which both drive and are driven by their activity. Optical imaging techniques are optimal for monitoring cellular responses from large neuronal populations; but they do not provide the temporal resolution required for the simultaneous detection and characterization of network oscillations. The technological gap in directly monitoring large numbers of identified neuronal types and simultaneously recording fast-time scale responses generated by hippocampal networks remains a major outstanding challenge. In response to this unmet need, we have developed a novel ‘E-Cannula’ technology; a two-photon imaging window-cannula integrated with transparent graphene microelectrodes for simultaneous optical imaging and electrical recordings from the same neural populations across large areas.

Hippocampal sharp wave ripples (SWRs) are the most synchronous population pattern recorded in the mammalian brain^3^. They have been hypothesized to control the information transfer from the hippocampus to down-stream cortical structures for learning and memory consolidation as well as planning, credit-assignment, and prediction^3-10^. In this work, we applied the E-Cannula to obtain a dynamic map of SWRs in the CA1 region of the mouse hippocampus through simultaneous two-photon calcium imaging and direct recordings of SWR generation and propagation across the same population of cells. To date, *in vivo* studies into SWR-associated hippocampal neuronal dynamics have almost exclusively utilized electrophysiological recordings with single- or multiple-shank penetrating probes, with only a few instances of two-photon calcium imaging with simultaneous contralateral local field potential recordings^11,12^. Therefore, correlated spatial and temporal characteristics of SWR-related neural dynamics remain largely unknown. Furthermore, it is unclear how the activity of distinct neuronal assemblies relates to this untapped putative diversity in SWR events. To address these questions, we implanted the E-Cannula above the hippocampal area CA1 and performed simultaneous electrophysiological recording and two-photon calcium imaging of the intact CA1 region in head-fixed mice. We recorded both the multi-unit activity (MUA) and SWRs, while simultaneously monitoring fluorescence calcium signals from CA1 pyramidal neurons. We found that the MUA activity recorded by transparent microelectrodes increased during SWR events and was strongly phase-locked to nearby SWR events. SWRs often co-occurred with high synchrony events (HSE) detected with imaging, which involved coactivation of multiple neurons. We investigated the SWR activities recorded with transparent microelectrodes and found that SWR activity could be spatially local or global across CA1 region and could also exhibit stationary or travelling temporal characteristics. By performing decoding analysis, we found that the SWRs with different spatiotemporal patterns were associated with distinct patterns of underlying hippocampal neural activity. Finally, most hippocampal cells formed orthogonal cell assemblies whose activation was selective to SWRs displaying specific electric spatiotemporal patterns.

## Results

### *In vivo* multimodal recordings from hippocampal area CA1 with E-Cannula

In order to construct E-Cannula, we fabricated fully transparent graphene microelectrodes on flexible and transparent polyethylene terephthalate (PET) substrate (Figure 1A). We employed an electrochemical delamination graphene transfer method and a previously developed 4-step cleaning method^13^ for the fabrication (see Methods for details on the fabrication process). The transparent array consists of 16 100 μm diameter circular electrodes with 500 μm spacing. The scanning electron microscopic image shows the profile of the well-defined electrode openings (Figure 1A). The total area of the array was 2.3 mm × 1.8 mm to match the opening of the imaging cannula. The small size of the array was key to minimizing damage during implantation surgery. In order to reduce the overall array size without sacrificing the number of recording channels, we replaced the previous gold connecting wires with double-layer graphene wires and carefully routed the wires through the inner plane of the array instead of surrounding the array, significantly reducing the total size of the array by ∼4x times compared to the previous configuration^13^. We performed cyclic voltammetry and electrochemical impedance spectroscopy to characterize the graphene microelectrodes in 0.01 M phosphate-buffered saline (Figure 1B and 1C). The result from cyclic voltammetry measurements shows no redox peaks, suggesting that the electrode-electrolyte interface is dominated by the double-layer capacitance (Figure 1B). Our electrodes exhibited a uniform impedance across channels with an average value of 1.1 MΩ at 1 kHz, achieving 100% yield (Figure 1B).

**Figure 1.**
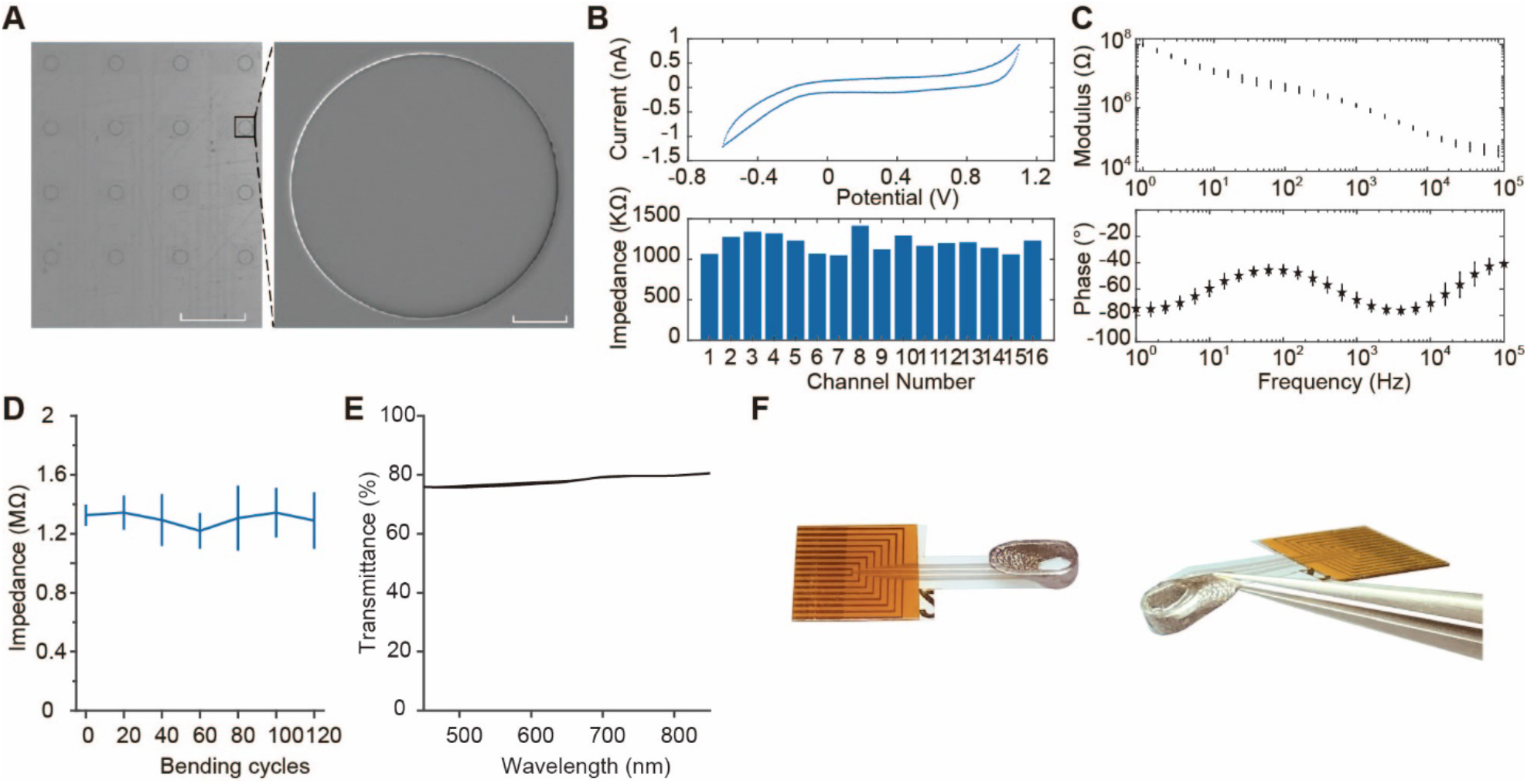
Fully transparent graphene microelectrode array and the integrated E-Cannula. **A**. Microscope image of the fully transparent 16-channel graphene microelectrode array (left) and the SEM image of one electrode (right). Left scale bar: 500 μm. Right scale bar: 20 μm. **B**. The cyclic voltammetry result of one example electrode (top) and the impedance of 16 microelectrode (bottom). **C**. The electrochemical impedance spectroscopy result showing the magnitude (top) and phase (bottom) of one example electrode. **D**. The result of bending test for the graphene array, showing stable impedance after at least 120 bending cycles. **E**. Transmittance of the graphene array under different wavelengths. **F**. Picture of the fully transparent array, the imaging cannula, and the integrated E-Cannula in top-view (left) and in side-view (right).

The flexibility and sturdiness of the transparent microelectrode arrays proved critical for their integration with the imaging cannula and their continued durability during chronic *in vivo* experiments. We characterized the reliability of the array with bending tests, where the array was bent to a radius of 5 mm. The results show that the impedance of the array stays stable even after 120 bending cycles (Figure 1D), demonstrating the high reliability of the array suitable for integration with the imaging cannula used in this study. Besides that, the array also exhibits high transparency across the wavelength ranges commonly used for optical imaging (Figure 1E). Figure 1F shows a picture of the fully transparent graphene microelectrode array, the imaging cannula, and the assembled E-Cannula. The flexible shank of the graphene microelectrode array was bent and attached to the outer wall of the imaging cannula and the fully transparent microelectrode array provided a completely clear field of view. In the present configuration the cannula was designed to have ∼30° angle. In practice, we found that the electrode impedance of the graphene array stayed roughly unchanged even after 90 degrees bending (Figure S1), making it compatible with cannulas of 90° angle thus further facilitating implantation.

With the fully transparent graphene microelectrode arrays, we performed simultaneous two-photon GCaMP-calcium imaging and electrophysiological recordings from the ipsilateral CA1 (Figure 2A, see Methods), while mice ran voluntarily and rested on a circular treadmill^14,15^. The E-Cannula was placed over the CA1 alveus of the hippocampus after aspiration of the underlying cortical tissues (see Methods), and 840×840μm images were acquired at 30Hz from CA1 stratum pyramidale (SP). As shown in Figure 2B and C, large populations of CA1 pyramidal cells were successfully imaged with single-cell resolution through the transparent graphene array (1079 ± 43, mean ± s.e.m. cells per field of view, FOV), and stably identified over 20 days after viral injections (Figure S2).

**Figure 2.**
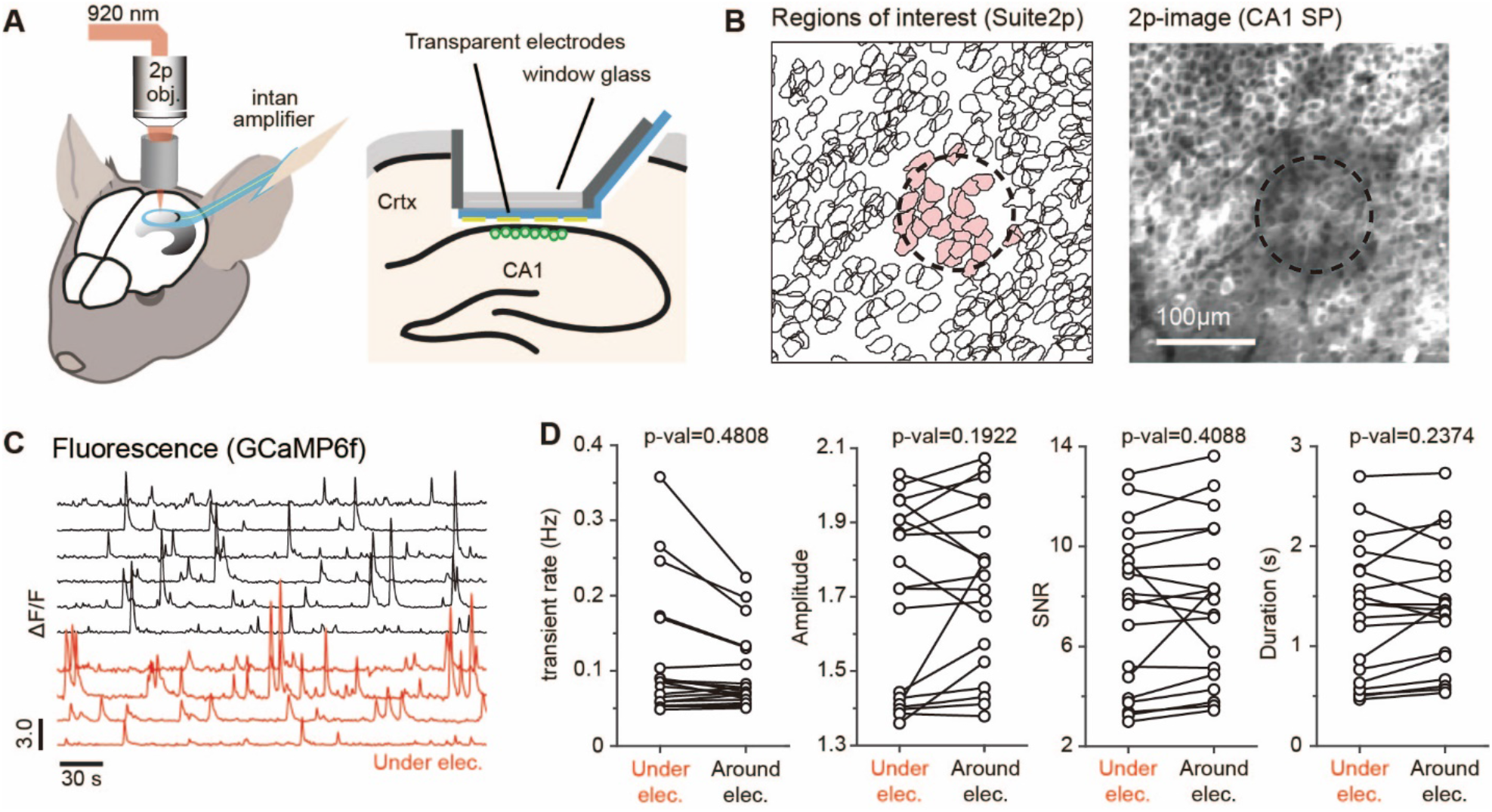
Simultaneous two-photon imaging and electrical recording from hippocampal CA1 area in awake mice. **A**. Schematics of in vivo multimodal recording from the dorsal CA1. The transparent graphene microelectrode array was attached to the bottom of canula and placed on the alvear surface of hippocampus. **B**. Example small area around a recording electrode within a larger imaging field-of-view (840 mm^2^) with imaging plane in the CA1 stratum pyramidale (*SP*). Dashed circle shows the location of the electrode. Scale bar = 100 mm. Left: suite2p-detected ROIs of CA1 pyramidal cells under (red shaded) and around the electrode. Right: time averaged two-photon image. **C**. Representative GCaMP calcium signals (ΔF/F_0_) extracted from cell ROIs around (black) and under (red) the recording electrode. **D**. Comparison of the transient rate, amplitude, SNR, and duration of GCaMP-calcium events between cells directly under the graphene electrode and around the graphene electrode, showing similar activity properties (two-tailed bootstrap test, 10,000 times, n=19 recording sessions from 3 mice).

To confirm that electrode attachment to cannula glass does not degrade the imaging quality, we compared the calcium signals imaged through E-cannula and standard cannula. Both signal properties are similar in signal-to-noise ratio (SNR), transient amplitude and duration (Figure S3). We also characterized the fluorescence activity between the cells under the graphene electrode versus around the graphene electrode and found similar amplitude, duration, event rate, and SNR of calcium events (Figure 2D). These results confirm that our E-cannula provide similar imaging quality compared to standard imaging cannula. Furthermore, unlike conventional metal-based electrode arrays, transparent graphene microelectrodes do not generate any light-induced artifacts that would otherwise interfere with electrophysiological recordings ^13,16^. Photo-induced currents are intrinsically very weak and fast in graphene, requiring special structures or extremely low temperatures to even detect them^17,18^.

We next examined features of local field potential (LFP) recorded from the array. We observed clear theta (4-8 Hz) oscillations in the recordings when the animal was running (Figure S4A and B). Consistent with the previous reports ^19^, we also observed phase-amplitude coupling in delta-gamma and theta-gamma bands (Figure S4C), indicating that CA1 strata pyramidale and stratum oriens were the main sources for the recorded LFPs. In a previous study, theta waves were reported to travel along the septotemporal axis of the hippocampus^20,21^. In our data, we also observed that the theta oscillations traveled along the septotemporal axis in the 2D recording space with a speed of 0.1484 m/s (Figure S4D and E), which is similar to the results reported in the previous study^20^.

We then employed E-Cannula to record SWRs across a wide area of dorsal CA1. Representative waveforms and the power spectrograms of detected SWRs are shown in Figure 3A. In agreement with previous electrophysiological and optical recordings^3,11,12,22^, we clearly observed that a large number of CA1 pyramidal cells were sharply activated during the SWRs detected in the channels within FOVs (Figure 3B and C). A significant fraction of synchronous calcium events (SCEs) co-occurred with SWRs (51% SWRs to SCEs, and 35% SCEs to SWRs, Figure 3D), and SCEs were associated with a robust increase in ripple band envelope power (Figure 3E). Together, the E-Cannula design allows us to simultaneously monitor calcium dynamics in large populations of individually identified cells and to relate these to the prominent oscillatory electrical potentials directly recorded from imaging FOVs.

**Figure 3.**
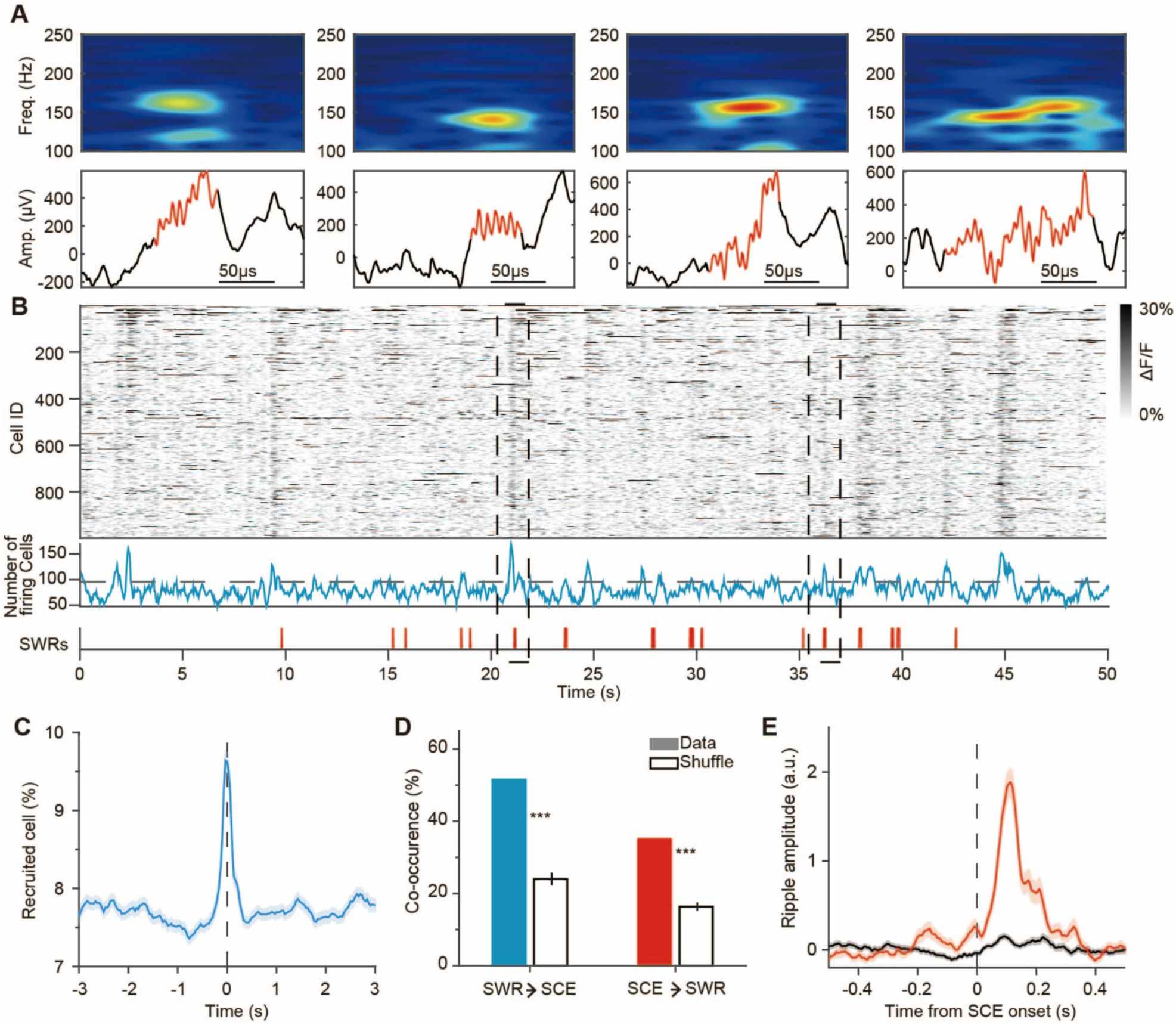
Synchronous dynamics of CA1 pyramidal cells during SWRs. **A**. Examples of SWRs detected on the electrode (top: ripple band power spectrogram, 100 to 250 Hz; bottom: LFP traces). **B**. SWR-associated SCE of CA1 pyramidal cells. ΔF/F of the cells during immobile states (top), the number of the active cells over time frames (middle). Black dash line indicates significance threshold for SCE detection. The onsets of detected SWRs on the electrode above FOV (bottom). Black dash lines indicate co-occurrence events of SCEs and SWRs. **C**. Peri-stimulus time histogram of percentage of firing pyramidal cells around local SWR onsets. **D**. The co-occurrence rate between SWRs and SCEs. The error bars indicate the s.e.m. \*\*\**P* < 0.001. **E**. Peri-stimulus time histogram of z-scored SWR envelop detected by the electrode above FOV around SCEs and non-SCEs.

### Multi-unit activities recorded by E-Cannula from hippocampus

Since previous studies have demonstrated that the spiking activity of hippocampal neurons could be detected at distance of up to 150-200 μm^23-25^, we examined whether our transparent graphene microelectrodes could also detect spiking activity from the hippocampus. Figure 4A shows an example signal after high-pass filtering at 500 Hz. Using Kilosort (see Methods), spikes were detected from multiple-channels, and sorted into discrete MUA clusters (number of clusters per mouse: 2.25 ± 0.75, firing rate: 0.77 ± 0.32 Hz, mean ± s.e.m.) that exhibited stable SNR across multiple days after implantation (Figure 4B). The spikes of each MUA were typically detected in one of the channels (Figure 4C) and stably exhibited across multiple days (Figure 4D). Detected spike waveforms showed both symmetric and asymmetric shapes with most MUA clusters exhibiting low firing rate (Figure S5A), suggesting that the detected MUA clusters primarily originated from pyramidal cells^26,27^. We further sought to determine whether spike timing of MUA clusters was modulated by SWRs. Consistent with previous findings that SWRs are associated with robust increase in CA1 neuronal firing rate^28^, we found that all the clusters clearly increased their firing rates around the onset of the SWR events detected on the same channel (Figure 5A). The MUA spikes were phase-locked to SWRs, predominantly occurring at rising edge of the ripple oscillation (Figure 5B), while the ripple-frequency phase-modulation of MUA gradually attenuated as the distance to the ripple channel increased (Figure 5C). Therefore, MUA clusters detected in the graphene array mainly included action potentials from neurons proximal to each channel, and phase-modulation by SWR could operate at local circuit level. Finally, we checked the simultaneously imaged calcium ΔF/F signals from the cells right below one recording channel and compared them with spikes detected from the same channel (Figure S5A). We computed the overlapping ratio of MUA spikes with the calcium events from individual cells and performed shuffling test. The result does not show significant matching between the MUA spikes and the cells (Figure S5B). Together, these results demonstrate that E-Cannula facilitates simultaneous detection of electrical and optical signals from hippocampal cells.

**Figure 4.**
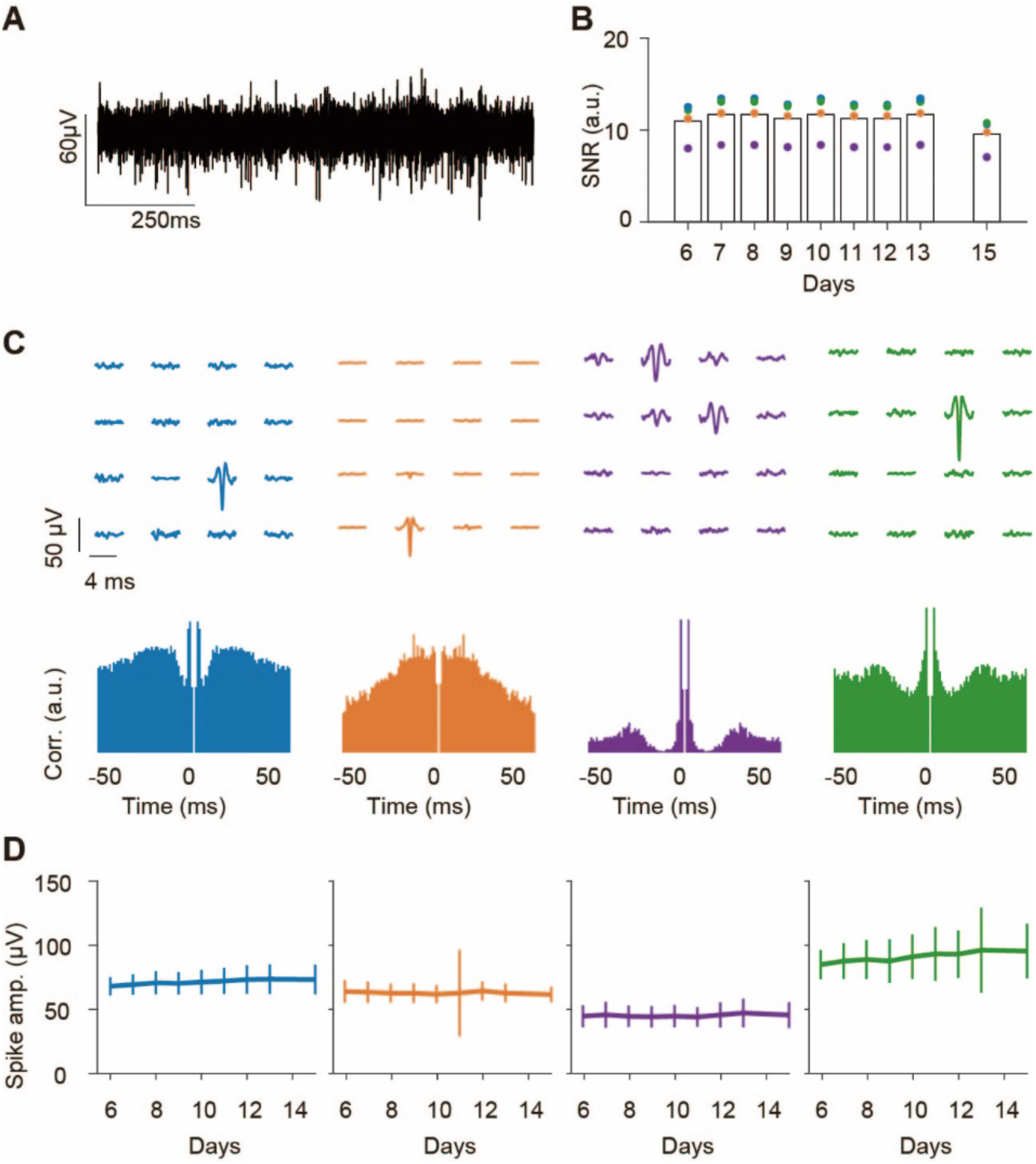
Hippocampal putative spikes recorded by the 2D graphene array. **A**. Representative LFP trace from one electrode after high-pass filtering at 500 Hz. **B**. SNR of the detected spikes at different days. Those detected spike cluster have stable SNR across days. **C**. Spatial profiles of the mean waveforms for each spike cluster (top) and the autocorrelograms (bottom). The spike waveforms were mainly detected on a single channel. **D**. Spike amplitude of each cluster at different days after implantation, showing a stable recording for putative spikes across days.

**Figure 5.**
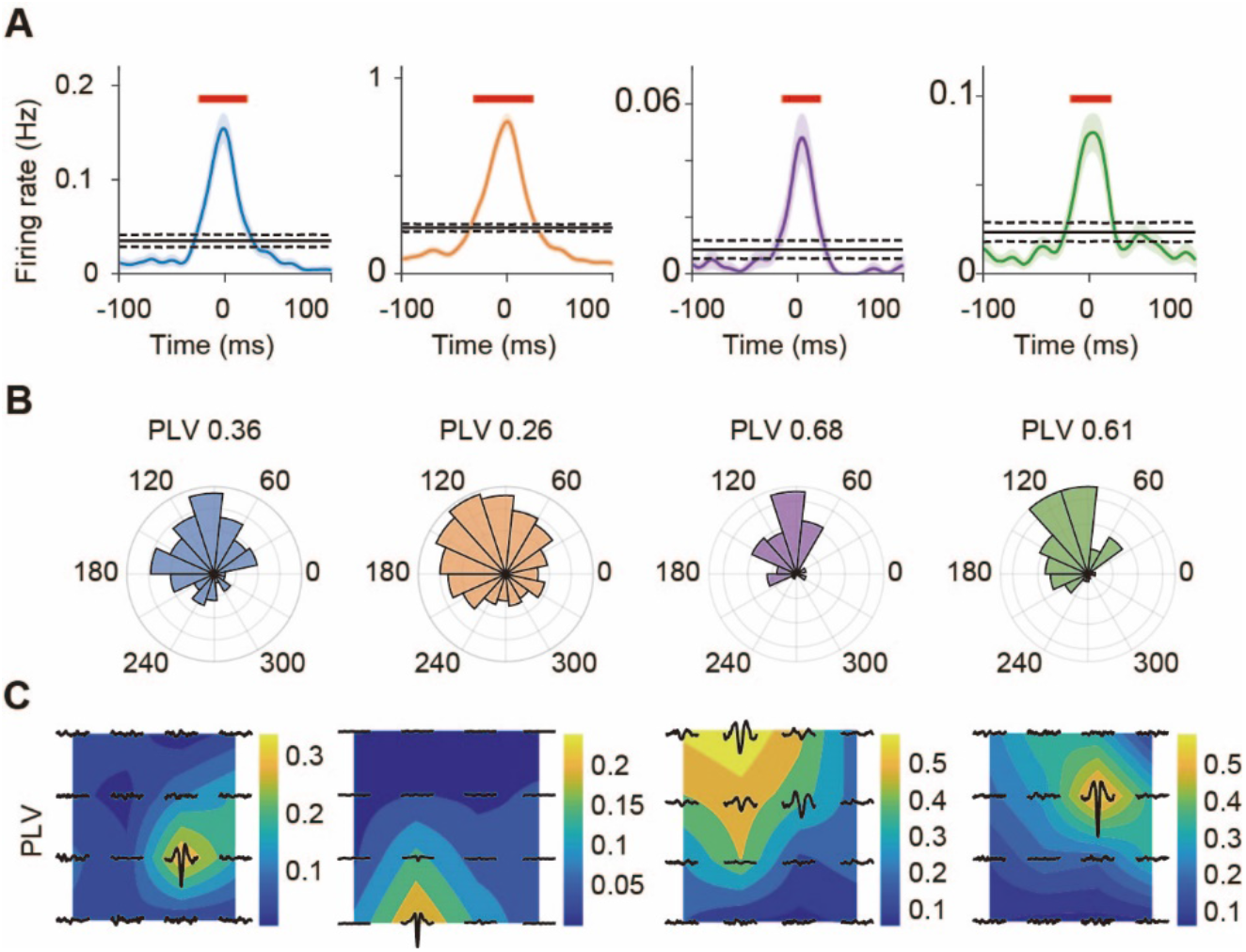
Locally controlled phase modulation of MUA spikes by SWRs. **A**. Firing rates of each spike cluster shown in *Figure 4* during SWR events detected on the same channel. All the spike clusters increase their firing rates around the SWR onset. **B**. The ripple band phase distribution at the putative spike firing times. All the spikes are phase-locked to the falling edge of the ripple waveforms. **C**. The phase locking values (PLVs) between the firing times of each spike cluster and the phase of SWRs at different electrodes. The putative spikes were mainly phase-locked to the SWR signals from the adjacent electrodes.

### Spatiotemporal characteristics of SWR events and their correspondence with hippocampal neuronal activity

Given that SWRs are generated by transient neuronal discharges propagating across the hippocampal circuit^28-30^, we hypothesized that SWRs could be biased by anatomical extent along the septotemporal and transverse axes, rather than being universally synchronized. To test this hypothesis, we first computed the power and peak latency of recorded SWRs for each channel relative to all the channels across the array. Then we carried out K-means clustering to identify clusters with distinct spatiotemporal patterns (see Methods). We found multiple SWR clusters characterized by different spatial occurrence and temporal propagation mode, further classified into four groups (local-stationary, local-traveling, global-stationary, and global-traveling) as shown in Figure 6A. The ratio of SWR events assigned to different clusters across days after implantation is shown in Figure 6B. Across different mice (n=3), we observed similar SWR clusters showing various spatiotemporal patterns (Figure S6).

**Figure 6.**
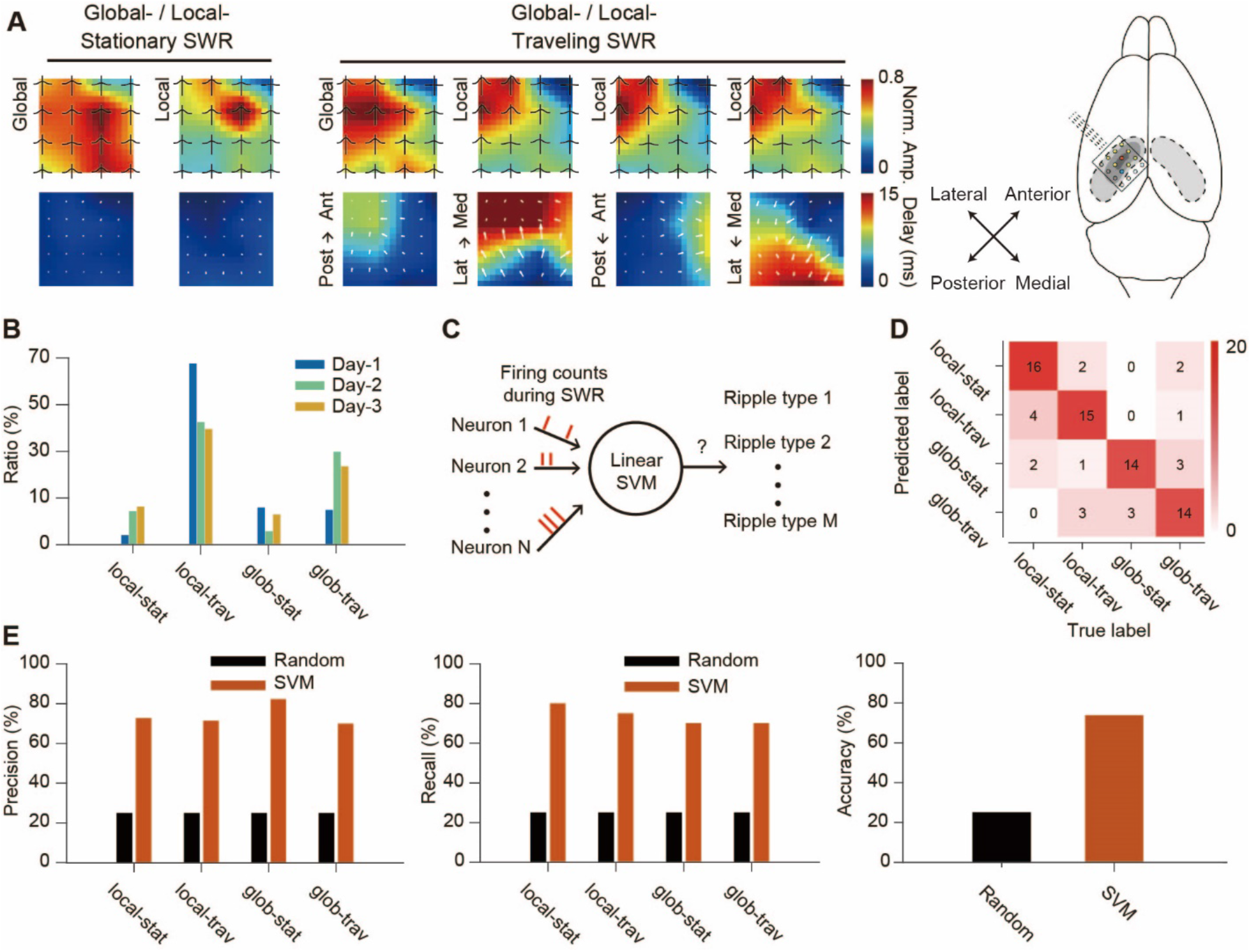
Spatiotemporal patterns of SWRs decoded by CA1 local circuit activity. **A**. Identified SWR clusters showing different spatial activation patterns (top) and temporal delays (bottom) relative to the septotemporal and transverse axes of the hippocampus. The black traces on the top row show the average ripple envelop detected at each channel. The white arrows on the bottom row show the numerical gradient (Med: medial; Lat: lateral; Ant: anterior; Post: posterior). **B**. Ratio of SWR events with different spatiotemporal characteristics across recording days. **C**. Decoding of SWR types based on cellular calcium signals around SWR onsets. **D**. Confusion matrix for decoding the spatiotemporal characteristics of SWRs. **E**. Precision, recall, and accuracy for decoding the spatiotemporal characteristics of SWRs.

We next sought to characterize neuronal activity patterns associated with SWR clusters with different spatiotemporal characteristics along the septotemporal and transverse axes of the hippocampus. For this, we applied decoding analysis with support vector machines (SVM) to ask whether the four groups of SWR events could be discriminated based on the neuronal activities (Figure 6C and see Methods). We used the calcium signals of the imaged cells during SWR as the input features and decoded the group identity of SWR event and performed recursive feature elimination to select the best subset of neurons that are most informative about the SWR cluster type. All the SWR clusters were successfully decoded with high accuracy, precision and recall above chance levels (Figure 6D and E and Figure S6). These results suggest that the spatiotemporal bias of SWRs are a potential mechanism for the selective activation of specific hippocampal cell assemblies.

### Differential recruitment of cell assemblies to SWR clusters

Having seen that the SWR cluster identity could be decoded from the calcium activity dynamics of hippocampal neuronal populations, we asked whether individual neurons formed into cell assemblies that exhibited segregated activities under different SWR clusters. For this, we computed the similarity between the calcium activity of recruited cells during each SWR event and performed clustering with community detection to assign the cells into different groups (see Methods). We identified multiple cell assemblies with size ranging from 30 to 140 cells (Figure 7A), the topological distribution of the cell assemblies was unbiased with respect to the anatomical coordinates of cells within the FOVs (Figure 7B and Figure S7). To examine the associations between each cell assembly and the SWR clusters, we analyzed the firing ratio of the cell assemblies during each SWR cluster. Notably, we found that all the cell assemblies exhibited selective firing that were higher or lower than average for specific SWR clusters, and that the preferred cluster identity varied across assemblies (Figure 7C-E). We repeated this analysis for all the mice yielding similar results (Figure S8). Thus, our results demonstrate that E-Cannula recordings allow for the identification of cell assemblies that are differentially recruited to SWR clusters with distinct spatiotemporal profiles along the major axes of the hippocampal CA1.

**Figure 7.**
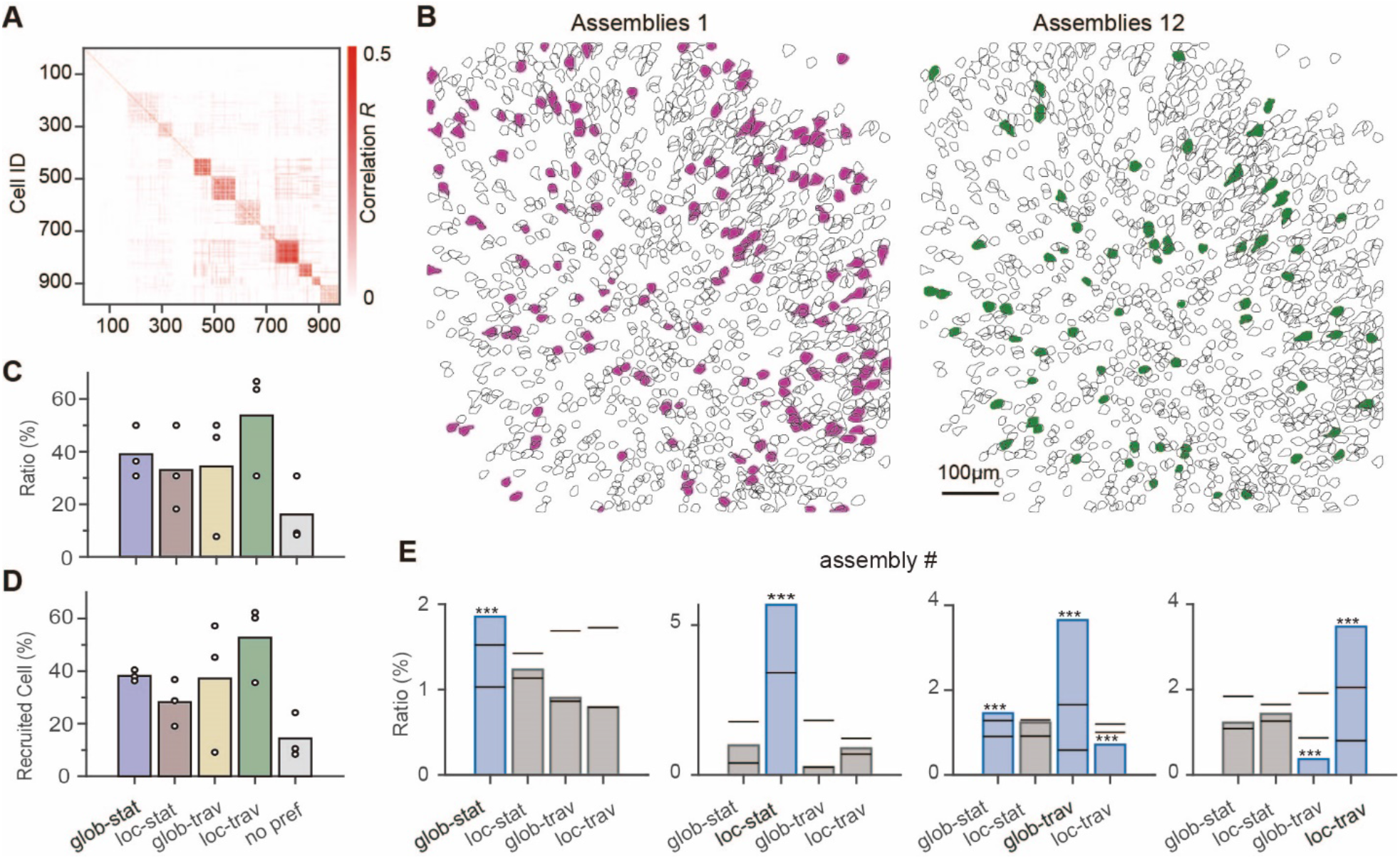
CA1 cell assemblies differently activated during spatiotemporal-profiled SWRs. **A**. Adjacency matrix of the identified cell assemblies. **B**. Topological distribution of cells for example cell assemblies. **C**. Ratio of cell assemblies associated with SWR events of different spatiotemporal characteristics. **D**. Ratio of recruited cells from assemblies associated with SWR events of different spatiotemporal characteristics. **E**. The firing ratio of cell assemblies under different SWR clusters. All cell assemblies show significant higher/lower participation ratio under different clusters. The ratio of each cell assembly under each SWR cluster was tested by shuffling test (P < 0.05 with FDR correction, \**P* < 0.05, \*\**P* < 0.01, \*\*\**P* < 0.001). The bars are the 2.5 percentile and 97.5 percentile values obtained from shuffled data.

## Discussion

In this study, we report the development and implementation of E-Cannula technology for simultaneous *in vivo* electrophysiological recordings and two-photon calcium imaging from the same hippocampal tissue volume. In vivo recordings of the hippocampal dynamics have mainly utilized technologies with single modality. With our E-Cannula, we demonstrate reliable multimodal recordings of MUA spikes, SWRs, and cellular calcium activity dynamics, combined simultaneously in the same experiment. This combination of multiple modalities allowed us to carry out a comprehensive characterization of SWRs across spatial and temporal resolutions, providing a more detailed and complete picture of SWR-related spatiotemporal neural dynamics that was inaccessible for conventional single modality approaches.

We find that the SWR activity across 2D space exhibits diverse spatiotemporal patterns, including different propagation directions and spatial spread. These results are consistent with previous studies using penetrating electrode array with multiple shanks spanning in 1D direction^31^. Due to limitations on implanted electrode arrays, these reports were able to observe traveling SWR activities only along the antero-posterior axis. In our experiment, we confirmed the traveling behavior of SWR activities along this main axis, but further report that SWRs exhibit variable spatial extent along multiple axes in 2D space.

While E-Cannula allowed us to detect MUAs, unsurprisingly we did not observe high level of correspondence between the putative spikes and calcium activity of neurons under the recording channel. The ability of our approach to register putative spikes recorded by the transparent electrodes to individual cells imaged directly beneath the electrode could be constrained by various factors. First, the calcium response may not reflect all of spiking events of the cell, leading to potential false negative responses. Previous studies have shown that the fluorescence activity could be silent during sparse firing events or under low magnification of the imaging^32^. Second, considering the electrode size and the distance to the pyramidal layer and the stratum oriens of the CA1 region, the detected MUA spikes likely originate from multiple pyramidal cells and potentially also from interneurons, further limiting reliable mapping of the detected spikes to imaging signals.

Recent studies have successfully combined local field potential recordings from the hippocampus from one hemisphere with simultaneous two-photon calcium imaging from the other hemisphere^11,12^. They reported that the SCEs in the hippocampus co-occurred with the SWR events detected in the contralateral hippocampus. Our E-Cannula technology allowed simultaneous recordings of the electrical and fluorescence activity from the same hippocampal neuron population. In our data, we observed higher co-occurrence ratio between the synchronous calcium events and SWRs, revealing a more direct correspondence between the optically detected population events and the SWRs. Besides the co-occurrence ratio, a previous study also reported that different cell assemblies were recruited during synchronous calcium events. Our E-Cannula approach was able to replicate these findings and further demonstrate that these cell assemblies are differently recruited during SWR cluster with distinct spatiotemporal characteristics. Notably, the recruitment of unique cell assemblies by SWRs with distinct spatiotemporal patterns provides a potential mechanism for the orthogonal reactivation of assemblies associated with discrete memories while minimizing memory interference.

To the best of our knowledge, our study is the first to demonstrate a successful combination of simultaneous 2D electrical recordings and two-photon calcium imaging from a large area in the hippocampus. In the future, the spatial resolution of electrical recordings by E-cannula could be further enhanced by reducing both the size of the graphene electrode and the spacing between adjacent recording channels, enabling the investigation of hippocampal CA1 neural circuits at a finer scale. Also, shrinking the electrode size and pitch will improve the spike recording precision and electrical source separation of the E-cannula, making it possible to simultaneously record calcium signals and action potentials from the same neurons *in vivo*. Future work could further leverage the temporal-resolution of E-cannula electrical recordings with the cellular and genetic targeting techniques utilized in calcium and voltage imaging to dissect the interaction between the activity of specific cellular motifs and anatomically heterogeneous network oscillations. Moreover, the demonstrated electrical recording and imaging stability of the E-cannula is ideally suited for future comprehensive longitudinal tracking of the cellular and regional network changes underlying long-term memory. Finally, the above mentioned advantages of the E-cannula system make it well suited for advancing research on the mechanistic relationships between local and global markers of activity across time. This research in turn may have implications in the development of robust brain-machine interfaces. Therefore, our E-Cannula approach and associated experimental protocols has the potential to advance the scope of future chronic recording experiments, including facilitating the dissection of the large-scale hippocampal activity dynamics which underlie learning and memory consolidation.

## Methods Summary

### Graphene transfer and four-step graphene cleaning protocol

The electrochemical delamination transfer method was adopted to achieve less poly(methyl methacrylate) (PMMA) residue on the surface of the transferred graphene^13,33^. 300 nm thick PMMA (Microchem, 495 PMMA A4) was spin coated on chemical vapor deposition (CVD) grown monolayer graphene on copper film (GROLLTEX). In PMMA/graphene/copper layers, copper was connected to the cathode of the DC power supply and the anode was dipped into 0.05M NaOH. With applying 20V DC bias, PMMA/graphene/copper layers were gradually immersed into the NaOH solution with forming hydrogen gas bubbles induced between the graphene and copper layer, which delaminated the PMMA/graphene layers from the copper film. To remove the NaOH residues, the PMMA/graphene layers were transferred to deionized water 3 times. Then, PMMA/graphene layers were transferred on the polyethylene terephthalate (PET) substrate and dried at room temperature. After the PMMA/graphene layers were fully dried, the sample was annealed at 125°C for 5 min to improve graphene/PET adhesion and mitigate the PMMA wrinkles. To remove the PMMA layer, the sample was immersed in acetone for 20 min at room temperature, following by 10 cycles of Isopropyl alcohol (IPA) and deionized water baths (1 min for each cycle). To reduce the organic residues on the surface of graphene that affect the impedance of graphene electrodes and light-induced artifacts, a four-step cleaning protocol was adopted^13^. After graphene was patterned by using oxygen plasma etching, to remove the protection photoresist layer, the sample was immersed in i) AZ 1-Methyl-2-pyrrolidon (AZNMP) for 10 min and ii) Remover PG for 10 min to remove AZ1512 and PMGI, respectively. To cleanse AZNMP and remover PG, the sample was iii) soaked in acetone for 10min, following by iv) 10 cycles of IPA and deionized water baths. All the processes were conducted at room temperature.

### Double layer graphene microelectrode array fabrication

To achieve high optical transparency, we used 50 μm-thick PET as a substrate. 20 μm-thick PDMS was spin-coated on a 4-inch silicon wafer to provide mechanical support for PET and attached PET film on top of it. 10 nm Chromium and 100 nm gold were sputtered onto the PET substrate film using Denton Discovery 18 Sputter System. Metal wires and contact pad for zero insertion force (ZIF) connector were patterned with photolithography and wet-etching. The first graphene layer was transferred by using the bubbling transfer method as described above. After cleaning the first graphene layer, the second layer graphene was transferred and cleansed with the same procedure as the first layer graphene. To minimize the photoresist residue and protect the mechanical damage during patterning the double-layer graphene, PMGI/AZ1512 bilayer photoresist was applied on the double-layer graphene: i) 100 nm PMGI SF3 was spin-coated at 3000 rpm for 45 s and baked at 125°C for 5 min and ii) 1.2 μm AZ1512 was spin-coated at 4000 rpm for 45 s and baked at 95°C for 1 min. This bilayer photoresist was patterned with photolithography and double-layer graphene was etched with oxygen plasma etching (Plasma Etch PE100), following by a four-step graphene cleaning protocol as described above. To define the neural signal recording area, a 3-μm thick SU-8 encapsulation layer was spin-coated at 1500 rpm for 45s and baked at 95°C for 2 min. This encapsulation was patterned with photolithography and developed with SU-8 developer for 1min. To remove the SU-8 residue on the opening or recording graphene area, the sample was rinsed with 10 cycles of IPA and deionized water baths.

### Mice and viruses

For all experiments we used adult (8-16 weeks) male and female mice (two R1Ag5 transgenic mice, B6.Cg-Tg(Camk2a-cre)2Szi/J, The Jackson Laboratory, Jax #027310, and two wild-type mice). Mice were housed on a 12-h light dark cycle in groups of 2-5 mice, and individually after surgery for implantation. Recombinant adeno-associated viruses [rAAV1.Syn.FLEX.GCaMP6f.WPRE.SV4, Addgene #100833] to R1Ag5 mice or [pENN.AAV.CamKII.GCaMP6f.WPRE.SV40, Addgene #100834] to wild-type mice were used for imaging of pyramidal neurons.

All experiments were conducted in accordance with US National Institutes of Health guidelines and with the approval of the Columbia University Institutional Animal Care and Use Committee. No statistical methods were used to predetermine sample sizes. The experiments were not randomized, and the investigators were not blinded to allocation during experiments and outcome assessment.

### Surgical procedures

Stereotaxic rAAV injections were performed with a Nanoject syringe, as previously described ^34^. Mice were anesthetized with isoflurane and treated with 0.1 mg/kg buprenorphine or meloxicam for analgesia and 180nL (60nL per spot) of diluted virus (1:2 in sterile cortex buffer) was injected into left dorsal hippocampal CA1 (from Bregma AP -2.2, ML -1.75, and DV -1.0, -1.1, and -1.2 mm). E-Cannula was constructed by a graphene microelectrode, a 3-mm diameter cover glass (64-0720, Warner), and a stainless-steel cannula (bottom face: 3-mm diameter circle, 1.5-mm height, 45-angle of inclination at one side). The cover glass was first attached to cannula with optical adhesive (Norland optical adhesive 81). Then we applied a thin layer of optical adhesive to the surface of the cover glass and attached the graphene microelectrode array. The shank of the graphene microelectrode array was also attached to the wall of the cannula with optical adhesive. Finally, we used ultraviolet light to cure the adhesive until all the components were held in place. After 3-4 days for recovery, mice were implanted with E-Cannula over the left dorsal hippocampus as well as a steel head-bar for head-fixation. The scalp was removed, a 3.0 mm diameter craniotomy centered over the injection location was performed using a fin-tipped dental drill, and then the lateral side (∼1.0 mm at 2.35 rad) was sculpted to precisely match the size of E-Cannula base. The dura was removed, and the underlying cortex aspirated until fibers within the alveus were visible, while chilled cortex buffer was constantly irrigated. Stopping any residual bleeding with gel foam, we gently wedged E-Cannula into the craniotomy and lowered until it tightly touched the alveus fibers. After cannula was glued to skull, we bended electrode wire, attached to the cannula inclination, and covered all components with dental cement. A stainless-steel jewelers screw served as a ground/reference were inserted into the cerebellum.

### In vivo two-photon imaging and electrophysiological recording

After recovery from surgery, mice were handled for several days, and habituated to head-fixation and experimental setup. Mice were subsequently trained to run on the treadmill. Multimodal recording experiments started at 10 days following implantation surgery.

All imaging was conducted using a two-photon 8 kHz resonant scanner (Bruker) and 16x NIR water immersion objectives (Nikon, 0.8 NA, 3.0-mm working distance, respectively). We acquired 840 × 840 μm images (512 × 512 pixels) at 30 Hz using a 920-nm laser (50-100 mW, Chameleon Ultra II, Coherent) from CA1 stratum pyramidale. Green (GCaMP) fluorescence was detected with a GaAsP PMT (Hamamatsu Model 7422P-40). The preprocessing steps for acquired fluorescence signal using the SIMA software package were described in our previous work ^35^.

After SIMA-based motion correction, we detected neural regions of interest (ROIs), extracted associated fluorescence signals, and calculated neuropil-decontaminated ΔF/F_0_ using the Suite2p software package with built-in neuropil subtraction^36^. Identified ROIs were curated post hoc using the Suite2p graphical interface to exclude non-somatic components.

The microelectrode array of the E-cannula was connected to a customized connector board that routed the electrical signals to the Intan RHD2132 amplifier. Electrophysiological recordings were done with a multichannel recording system (Intan Technologies) synchronized with the AOD imaging system. The data was recorded at 20 kHz. In total, four mice were recorded.

### SWR detection and spike sorting

The detection of SWRs was performed by the following procedures^37^. The raw recording signals were first band-pass filtered between 120-250 Hz (4th order Butterworth filter) in both forward and reverse directions to prevent phase distortion. Then we applied Hilbert transform to extract the envelope of the ripple-band signals. To detect the potential SWR events in a single channel, we set a threshold to 4.5 standard deviations above the mean. When the ripple envelope crossed the threshold, a candidate SWR event was labeled and its start and end were further defined as the time when the ripple envelope returned back to the mean level. Similar to previous study, we merged the two candidate SWR events whose inter-event interval was less than 10 ms and discarded the candidate SWR events with duration less than 20 ms. After we identified the SWR events detected in each individual recording channel, we further defined the array-level SWR events. The start time was chosen as the earliest start time in all the channels detecting SWR, whereas the end time was chosen as the latest end time in all the channels detecting SWR. Finally, for all the automatically detected array-level SWR events, we further performed manual curation to discard the artifacts.

The spike sorting was performed with Kilosort 2^38^ to obtain the putative spike clustering results. Then we performed manual curation with Phy^39^ to refine the clustering results by checking the spike waveforms and the auto- and cross-correlograms. After rejecting the clusters corresponding to noise, we labeled the rest of the good clusters as multi-unit (MUA) since their spike waveforms resembled typical action potential waveforms, but the auto-correlogram exhibited a certain violation of the refractory period.

### Detection of synchronous calcium events and co-occurrence with SWR

To detect the synchronous calcium events (SCEs), we followed similar approach in previous study^12^ with slight modifications. After computing the deconvolved spikes of all the imaged cells, we used thresholds of 3 standard deviation to define the firing intervals for each cell. Then for each time frame, we count the number of firing cells in the adjacent 5 time frames (∼160 ms). To obtain the chance level firing cell number, we circularly shuffled the firing intervals 20 times for each cell and pooled the resulting firing cell number at all time frames. The 95 percentiles of the shuffled was chosen as the threshold. Then we applied this threshold to the number of firing cells at each frame in the original data to identify the time of SCEs. When computing the co-occurrence of the SCEs and the SWRs, we used a 200 ms time window. When a SWR and a SCE occur within 200 ms, they were labeled as co-occurring. To determine the significance of the SWR-to-SCE and SCE-to-SWR co-occurrence ratios, the timing of SWR were randomly permutated 1000 times and the p-values were computed as the percentage of resulting chance-level co-occurrences that were greater than the original co-occurrence.

### Clustering of SWR events

Before the clustering, we need to compute the spatiotemporal features of SWR events. To do that, we first computed the envelopes of the ripple band (120-250 Hz) signals from all the channels and computed the cross-correlation between each envelope and the mean envelope across all the 16 channels to determine the channel-wise time delay. For any channel, if the maximum cross-correlation is less than 0.5, we did not attempt to evaluate its time delay. These situations mainly happened when the ripple signals in that channel exhibited complex envelope so that the accurate determination of its timing is difficult. If the number of channels with evaluated time delay is less than 13, we discarded that event. Otherwise, we performed interpolations to fill in the time delays for the missing channels. To capture the spatial activation profile of the SWR events, we computed the magnitude of ripple band power at each channel and normalized it with the maximal value among all the channels. Finally, we computed the difference between the envelope in each channel and a standard gaussian curve using dynamic time warping and discarded the SWR events with complex envelope shapes (distance > 3.5 SD of ripple envelopes) and only focused on the SWR events with single dominant peak in their envelopes. These time delays and the normalized power magnitudes were then scaled to have same standard deviations and used as features to capture the spatiotemporal characteristics of the SWR events recorded in the 2D electrode grid.

With the above features, we performed K-means clustering to assign each SWR event to different ripple cluster types. To determine the proper number of clusters, we computed the within-cluster sum of squared distances under different number of clusters and chose the “elbow” point. We repeated the K-means clustering 10 times and chose the best clustering results. The template for each ripple cluster was obtained by averaging over all the SWR events assigned to the same cluster. Finally, we further grouped the identified ripple clusters into four bigger groups (stationary-local, stationary-global, traveling-local and traveling-global) depending on whether the maximum channel-wise time delays and the of the ripple templates. The ripple clusters with maximum channel-wise time delay greater than 5 ms were considered as traveling. Otherwise, they were considered as stationary. For the template of each ripple cluster, we defined each channel as active if its normalized power magnitude was greater than 0.6. The ripple clusters with more than 10 active channels were considered as global. Otherwise, they were considered as local.

### The algorithm for decoding the SWR events with different spatiotemporal characteristics

To decode the ripple cluster types using the cellular activity of hippocampal neurons, we used the support vector machine (SVM). The mean ΔF/F activity of each neuron during the [-150 ms, +150ms] around SWR onset were used as input features for the SVM algorithm. Since different ripple clusters have different number of SWR events, the resulting dataset is unbalanced. To overcome this difficulty, we created a new balanced dataset by subsampling the SWR events of different ripple clusters to match the number of SWR events in the smallest ripple cluster. Then the SVM model was trained on this new balanced dataset. To identify the most discriminative neurons and prevent overfitting, we performed recursive feature elimination^40^ with a step size of 50 neurons at a time. To measure the decoding performance, we evaluated the accuracy, precision, and recall using 10-fold cross-validation and compared these metrics against the results given by a random classifier.

### Detection of cell assembly and the firing ratio under different ripple clusters

To detect the cell assembly, we first computed the ΔF/F of cells between [-150 ms, +150ms] around each ripple onset. Then we applied a threshold of 30% on the mean ΔF/F of each cell to label it as active or silent during each SWR event. By computing the Jaccard similarity between each pair of neurons, we obtained the similarity matrix. To determine the chance level similarity, we construct a surrogate dataset by shuffling the identity of the active SWR events for each cell and recomputed the similarity matrix. By repeating this process for 200 times, we obtained a null distribution of the chance level similarity between each neuron pair. We then set any pairwise similarity below that threshold to zero. Using thresholded similarity matrix for both the original dataset and the surrogate dataset, we performed clustering using community detection algorithm to identify the putative cell assemblies. In the surrogate data, we found that many small-sized cell assemblies could emerge randomly. Therefore, we defined a threshold on the size of cell assembly as the 99th percentile of the cluster size obtained from the surrogate dataset. We then used this threshold to discard any putative cell assemblies in the original dataset whose sizes are smaller. The clusters that survive all these criteria were treated as identified cell assemblies.

### Statistical tests

All statistical analyses were performed in MATLAB. Statistical tests were two-tailed and significance was defined by p-value pre-set to 0.05. The error bars and shaded regions around line-plots represent ± standard error of the mean (s.e.m.) unless otherwise noted. All the statistical tests are described in the figure legends and each test was selected based on data distributions using histograms. Detailed statistical procedures are described in each sub-section of Methods. Multiple comparisons were corrected for by Benjamini-Hochberg corrections.

## Data availability

Data available on request from the authors.

## Code availability

Data available on request from the authors.

## Acknowledgements

We thank members of the Kuzum and Losonczy labs for helpful discussions. This research was supported by grants from the ONR (grant no. N00014161253 and N000142012405), the NSF (grant nos. ECCS-2024776 and ECCS-1752241) and the NIH (grant nos. R21 EY029466, R21 EB026180 and DP2 EB030992) to D.K. S.T. is supported by JSPS Overseas Fellowship, A.L. is supported by the National Institute of Mental Health (NIMH) 1R01MH124047, 1R01MH124867; National Institute of Neurological Disorders and Stroke (NINDS) 1U19NS104590 and 1U01NS115530, and the Kavli Foundation. Fabrication of the electrodes was performed at the San Diego Nanotechnology Infrastructure of UCSD, a member of the National Nanotechnology Coordinated Infrastructure, which is supported by the National Science Foundation (NSF; grant no. ECCS-1542148).

## Author contributions

This work was conceived by S.T., X.L., D.K. and A.L. J.H.K. performed microelectrode array fabrication and characterization. S.T. performed all animal experiments. X.L., S.T., and A.G. analyzed the collected data, with contributions from M.R., Y.L., D.K., A.G., and A.L. X.L., S.T., J.H.K., D.K. and A.L. wrote the manuscript and all the authors edited it.

## Ethics declarations

### Competing interests

The authors declare no competing financial interests.

---

**Figure S1.**
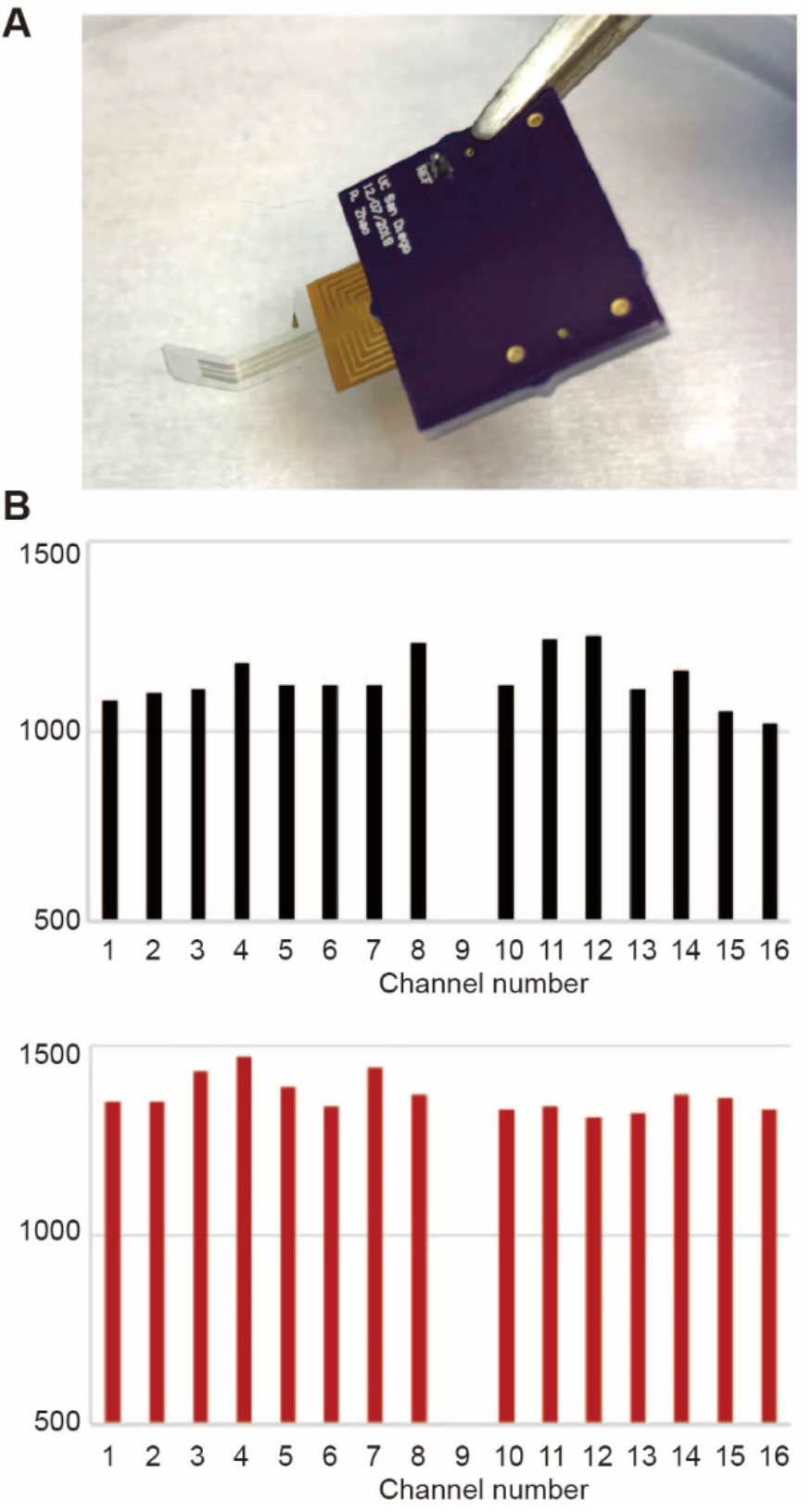
Electrode impedance after 90 degrees bending. **A**. Picture of the electrode array after 90 degrees bending. **B**. The electrode impedance before bending (top) and after bending (bottom). The electrode array still works with the impedance slightly increased after the excessive bending.

**Figure S2.**
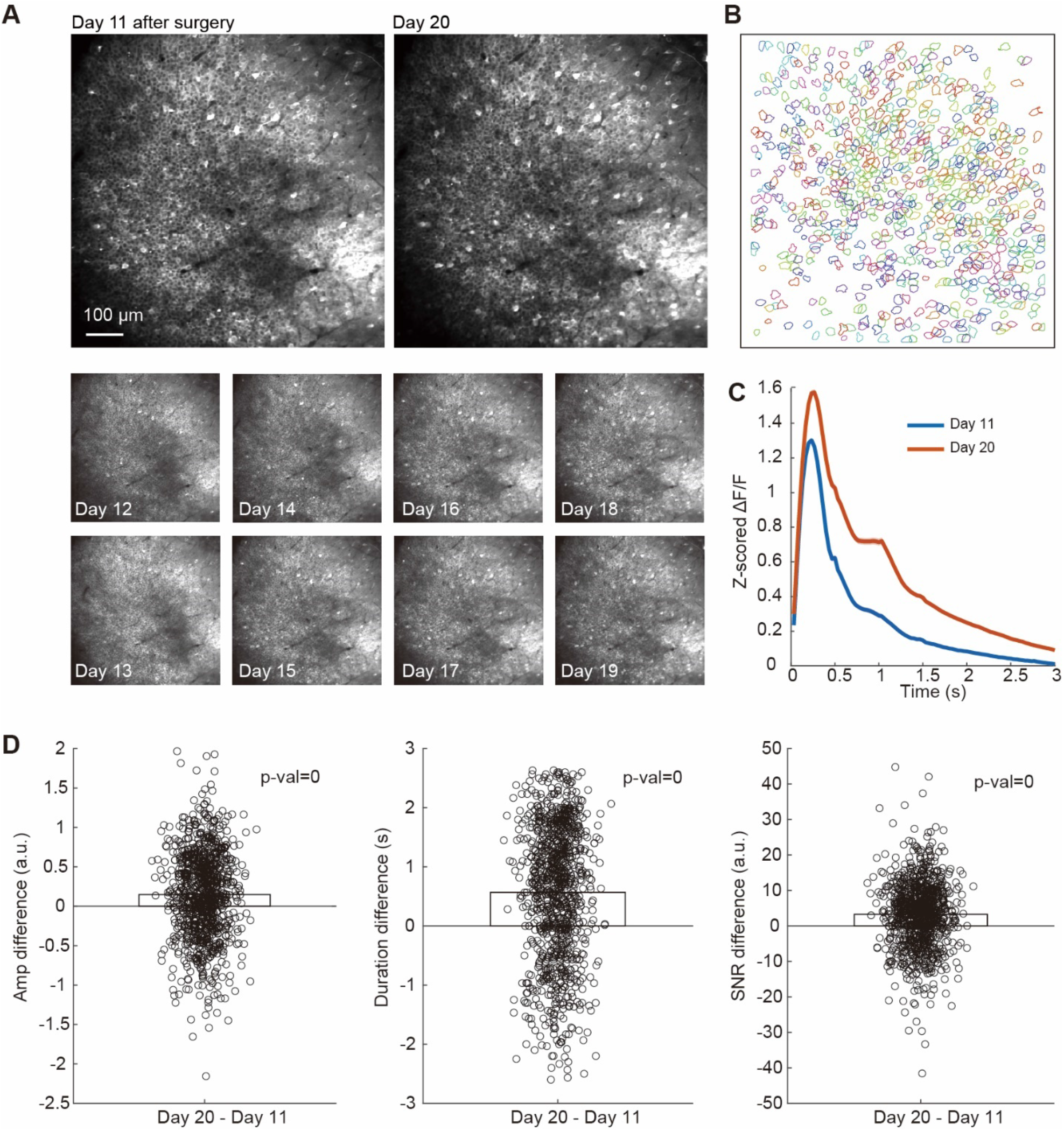
Comparison of imaging data obtained with E-cannula on Day 1 and Day 10. **A**. Example imaging field of view and region of interests obtained at Day 1 and Day 10. **B**. The SNR, amplitude, and duration of the ΔF/F response, showing stronger data response in Day 10 than Day 1. **C**. The mean ΔF/F response across cells on Day 11 (blue) than Day 1 (red). **D**. The comparison of the amplitude, duration, and SNR of GCaMP-calcium events in Day 11 and 20.

**Figure S3.**
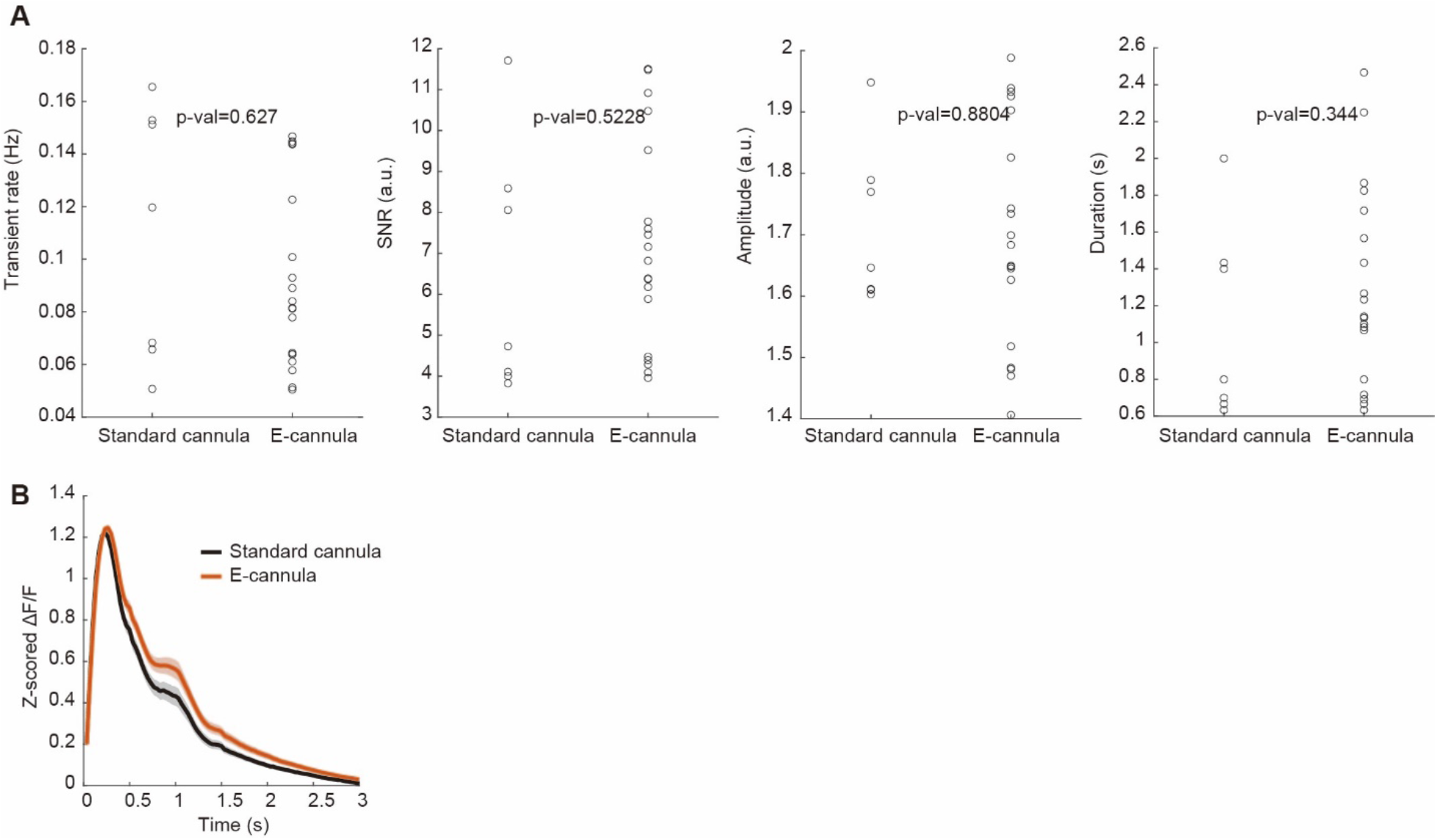
Comparison of imaging data obtained with standard cannula and the E-cannula. **A**. GCaMP-calcium transient rate, SNR, amplitude, and duration of the ΔF/F response, showing similar data quality between the standard cannula and the E-cannula. **B**. The mean ΔF/F response obtained with standard cannula and the E-cannula. The responses are similar (cluster-based permutation test).

**Figure S4.**
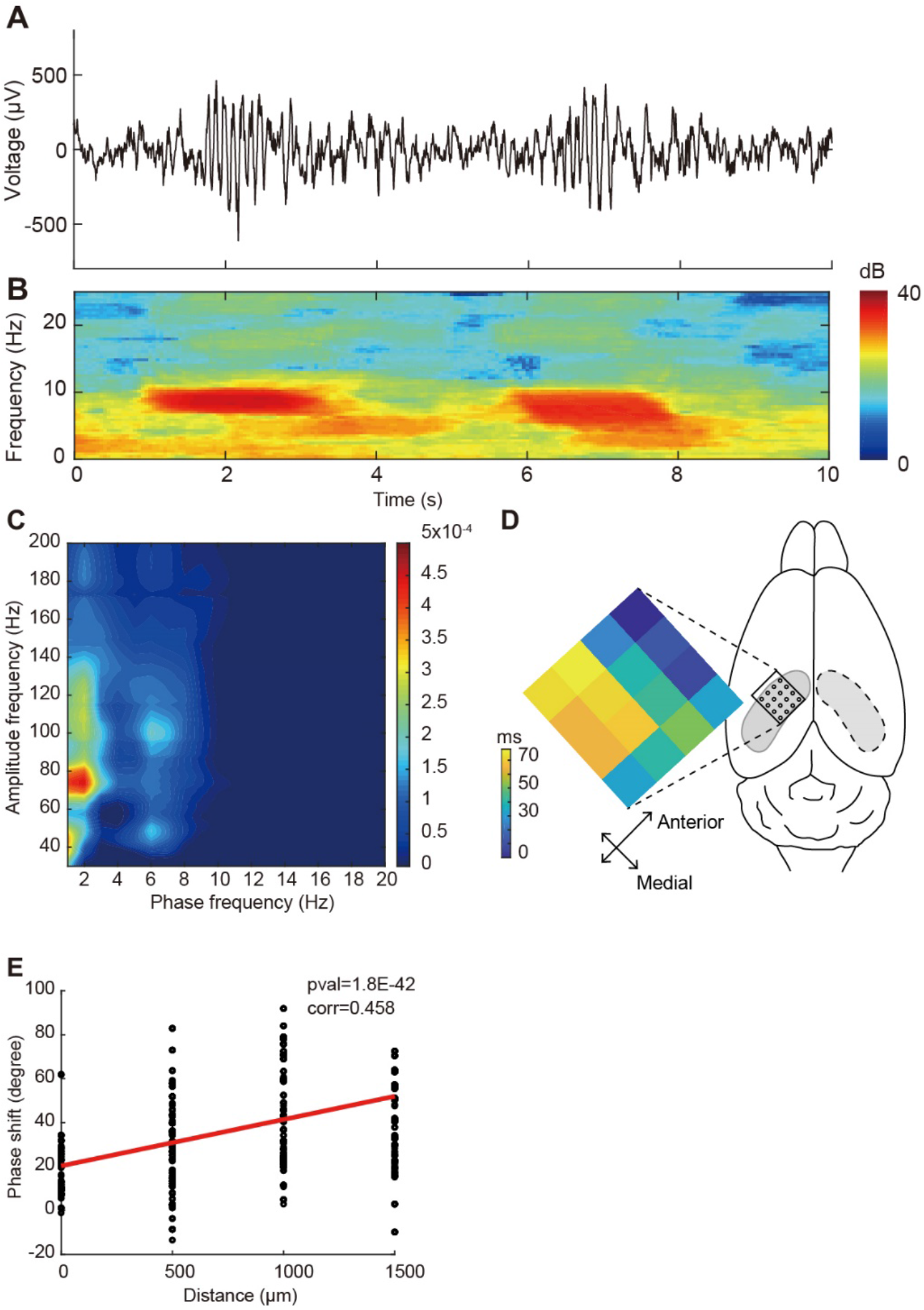
Characteristics of the theta oscillations detected by the E-cannula. **A**. Example LFP recordings from one channel showing two theta events. **B**. Spectrogram for recordings in a, showing power increase in theta band. **C**. Phase-amplitude coupling plot, showing significant theta-gamma and delta-gamma coupling in the electrical recordings. **D**. phase shift of theta band activity across array for one theta event (left) and the implantation map, showing obvious traveling of theta waves across the septotemporal axis. Corr = 0.458, P = 1.8E-42. Traveling speed = 0.148m/s. **E**. The phase shift of theta band activity as a function of septotemporal distance, showing a clear trend.

**Figure S5.**
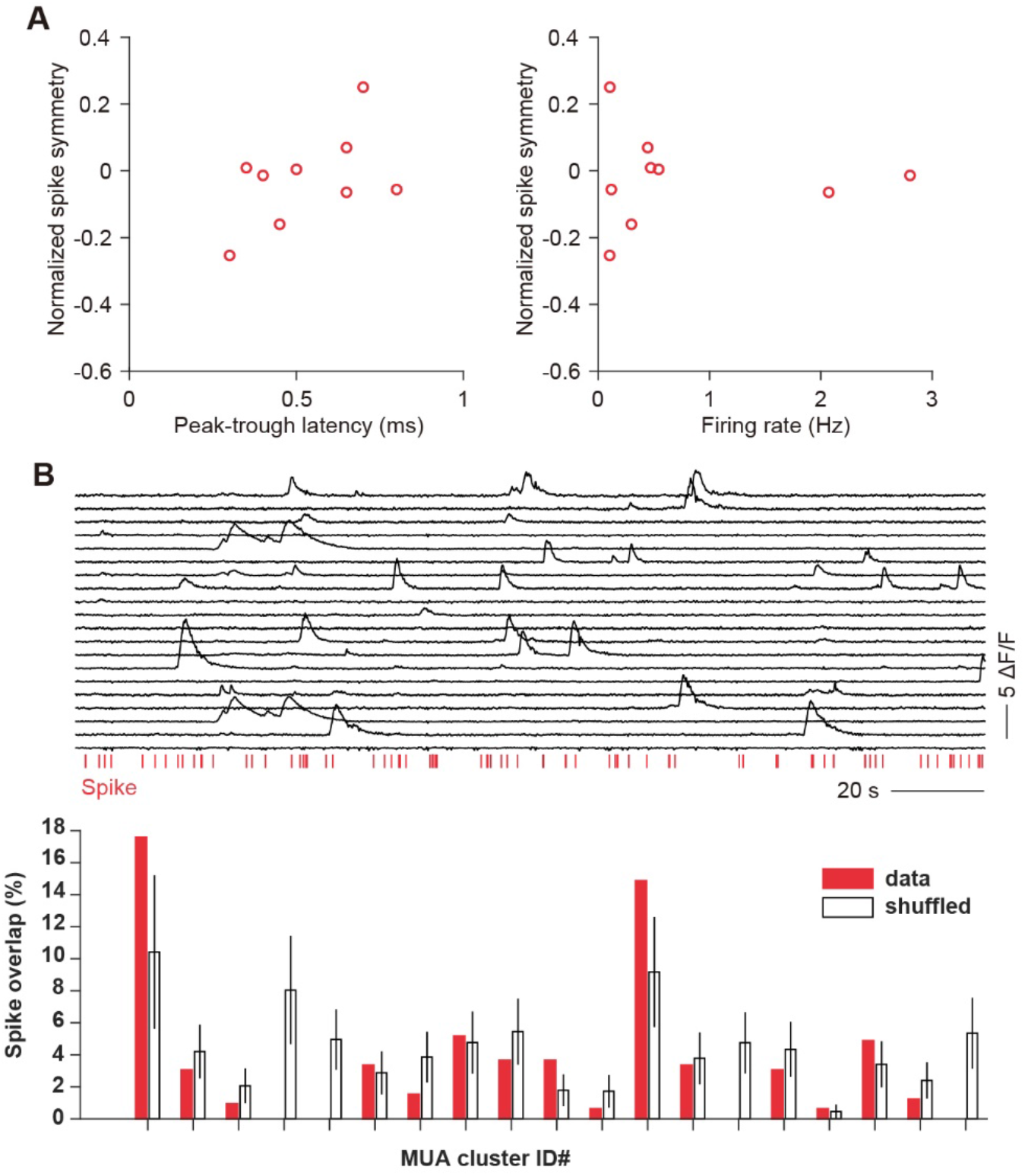
Waveform characteristics of MUA clusters and the simultaneously recorded fluorescence activity of cells. **A**. Each dot represents one detected MUA cluster. n=4 mice. ***B***. *Top*, the simultaneous recording of spikes from one spike cluster and the ΔF/F signals of 20 nearest cells right under the same recording electrode. Bottom, overlapping ratio of the recorded spikes with the calcium events from 20 cells.

**Figure S6.**
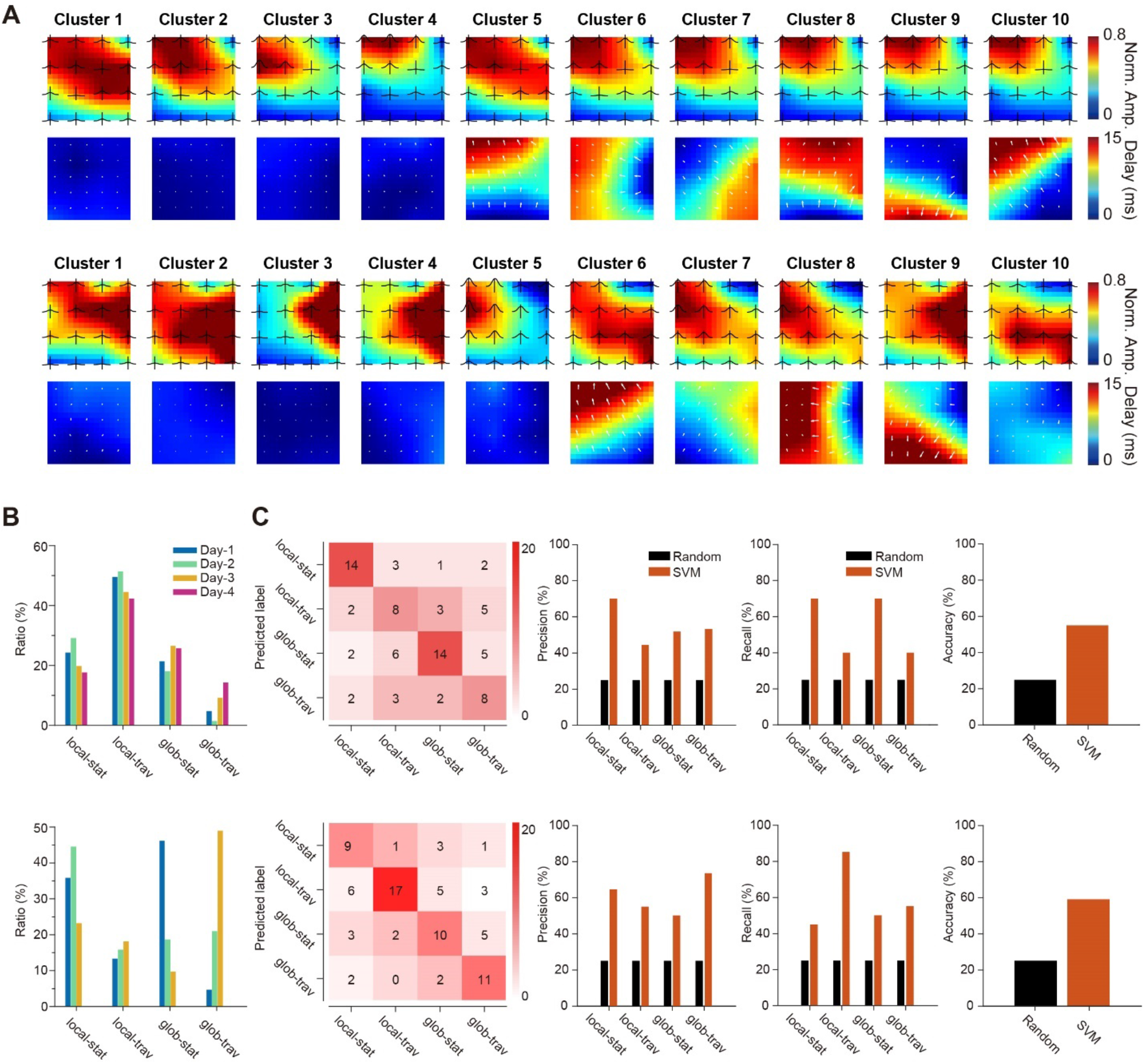
Spatiotemporal patterns of SWR events and its association to local cellular activity for the other two mice. **A**. Identified ripple clusters showing different spatial activation patterns (top) and temporal delays (bottom). **B**. The ratio of SWR events assigned to different ripple clusters in the recordings at different days. **C**. The confusion matrix and decoding performance (precision, recall, accuracy) for decoding ripple cluster types.

**Figure S7.**
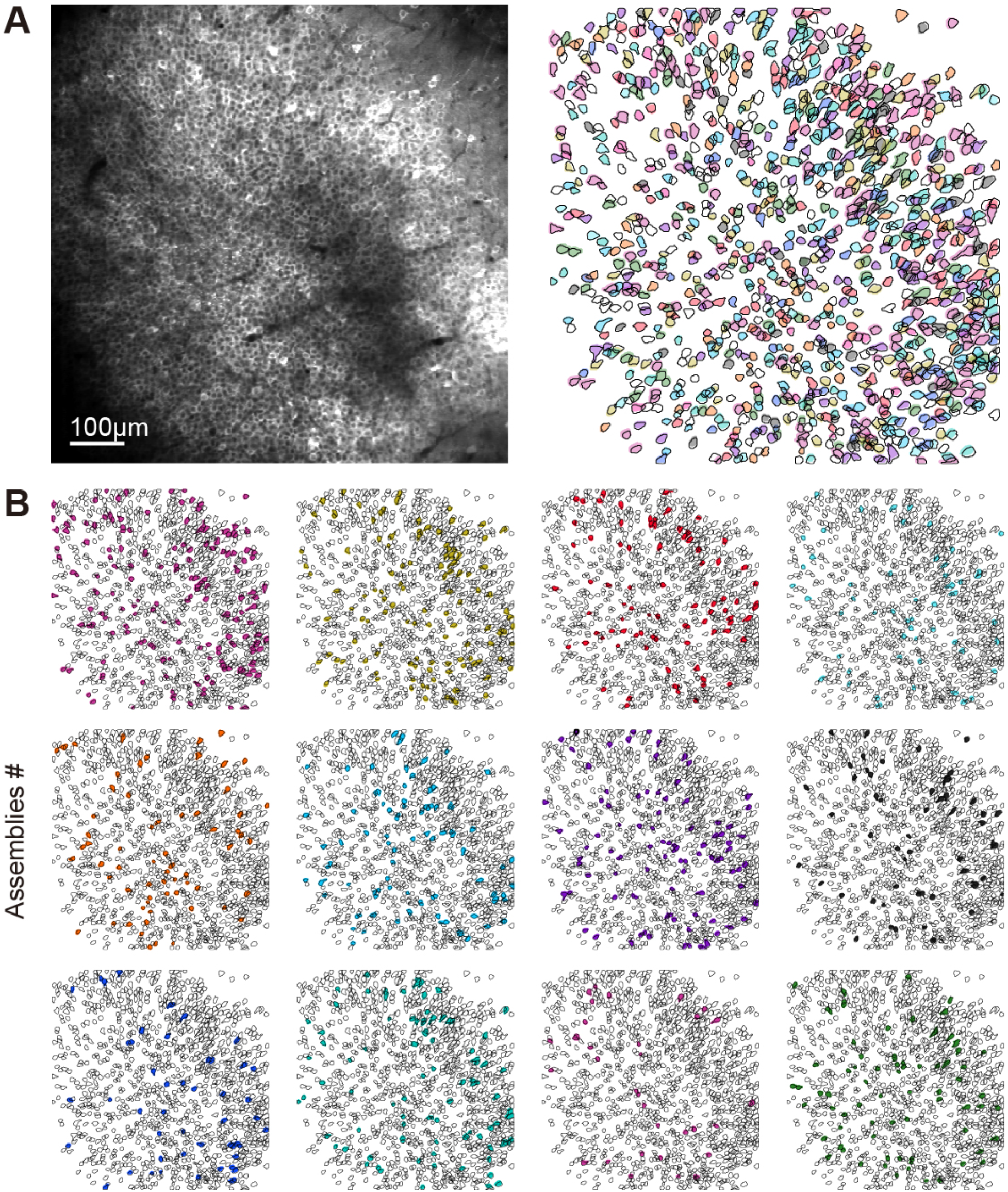
Additional data on topological distribution of cells within discrete cell assemblies. **A**. Representative FOV and ROI map of the imaged cells. Scale bar = 100 μmm. Filled colored ROIs illustrates the cells participating one of the assemblies, and each color indicates different assembly participations. **B**. all topological maps of the cells participating each assembly. The left top and right bottom maps are shown in *Figure 5B*.

**Figure S8.**
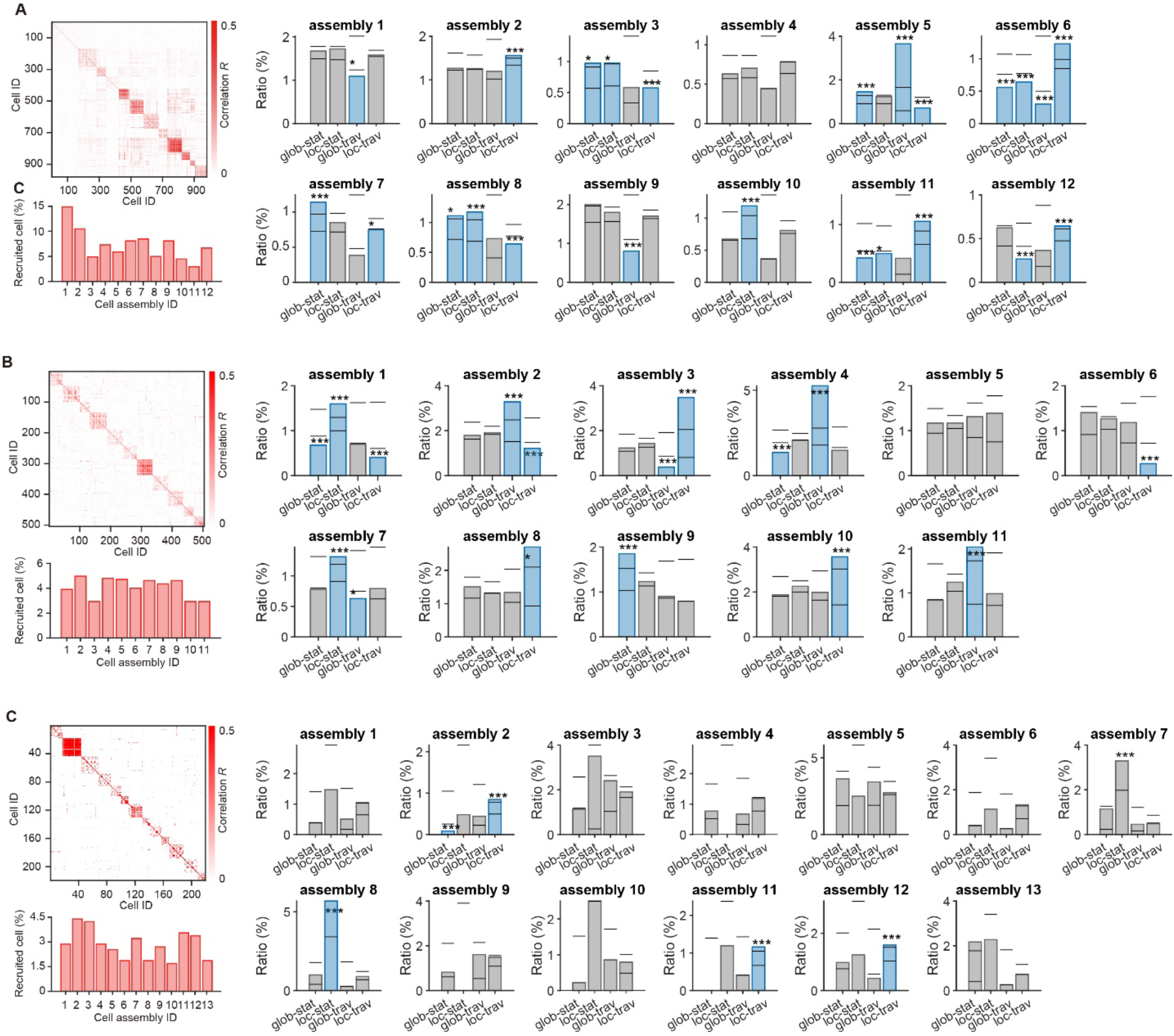
Activity of hippocampal cell assemblies during different ripple clusters for different mice. **A**. Adjacency matrix of the identified cell assemblies, the ratio of cells assigned to each assembly, and the firing ratio of cell assemblies under different ripple clusters. The ratio of each cell assembly under each ripple cluster was tested by shuffling test (P < 0.05 with FDR correction, \**P* < 0.05, \*\**P* < 0.01, \*\*\**P* < 0.001). The bars are the 2.5 percentile and 97.5 percentile values obtained from shuffled data. **B**. Same as in *(A)* with data from second mouse. **C**. Same as in *(A)* with data from third mouse.

## Notes

### Competing Interest Statement

The authors have declared no competing interest.

